# A high-throughput genome-integrated assay reveals spatial dependencies governing Tcf7l2 binding

**DOI:** 10.1101/2020.03.16.993204

**Authors:** Tomasz Szczesnik, Lendy Chu, Joshua W. K. Ho, Richard Sherwood

## Abstract

Predicting where transcription factors bind in the genome from their *in-vitro* DNA binding affinity is confounded by the large number of possible interactions with nearby transcription factors. To characterise the binding logic for the Wnt effector transcription factor Tcf7l2, we have developed a high-throughput screening platform in which thousands of 99-bp synthesised DNA sequences are inserted into a specific genomic locus through CRISPR/Cas9-based homology-directed repair, followed by measurement of Tcf7l2 binding by DamID. Using this platform at two genomic loci in mouse embryonic stem cells, we show that while the binding of Tcf7l2 closely follows the *in-vitro* motif binding strength and is influenced by local chromatin accessibility, it is also strongly affected by the surrounding 99-bp of sequence. The presence of nearby Oct4 and Klf4 motifs promote Tcf7l2 binding, particularly in the adjacent ~20 to 50-bp nearby and oscillating with a 10.8-bp phasing relative to these cofactor motifs, which matches the turn of a DNA helix. This novel high-throughput DamID assay provides a powerful platform to determine local DNA sequence grammars that causally influence transcription factor binding in controlled genomic contexts.

## 4 Introduction

Transcription factors recognise and bind to short DNA sequences, motifs, which can be measured directly through *in-vitro* binding assays, or discovered as enriched at sites bound across the genome. Such motifs, however, are insufficient to accurately predict where in the genome a transcription factor is bound, as most transcription factors bind to fewer than 10% of their strong motifs in any given cell type [Consortium, 2017]. Moreover, transcription factors exhibit cell-type specific binding patterns, despite no change in the DNA binding motif or genomic sequence. It is known that transcription factors influence each others’ binding, either through direct interactions, competition for binding sites, or indirectly by altering DNA organisation and accessibility. In different cell types it is then the set of transcription factors expressed that shapes their individual binding profiles. These interactions should be reflected in a “grammar”: a logic in how the organisation of individual transcription factor binding motifs shapes the higher order interactions.

The transcription factor Tcf7l2 provides a case study in inadequate prediction of cell-type specific binding patterns. Tcf7l2 is part of the Tcf / Lef transcription factor family (Tcf7, Tcf7l1, Tcf7l2, and Lef1) [Arce et al., 2006], which all bind DNA through a conserved high mobility group (HMG) domain that prefers the sequence SCTTTGWWS. This recognition occurs through through the DNA minor groove [Wetering et al., 1991, Wetering and C. Clevers, 1992], opening it up and creating a bend of 90-127 degrees [Love et al., 1995, Giese et al., 1995]. Tcfs act primarily as effectors of the Wnt signalling pathway, binding the transcriptional activator *β*-catenin, which upon Wnt signalling ceases to be constitutively degraded [Nelson, 2004].

As part of a conserved developmental pathway, Tcfs regulate various functions throughout different cell types in response to Wnt signaling. In mouse embryonic stem cells (mESCs) Tcfs appear necessary to reduce expression of other transcription factors that maintain pluripotency (notably Nanog) in order to allow differentiation [Pereira et al., 2006]. In the intestine Tcf7l2 helps maintain a constant proliferation of adult stem cells that support tissue renewal; a lack of dominant negative isoforms of Tcf7l2 and mutations in APC – part of the complex that enables GSK3*β* to cause degradation of *β*-catenin – is linked to colorectal cancers [Korinek, 1997]. Tcf7l2 knockout mice show problems with endoderm development and maintenance of intestinal stem cell populations [Korinek et al., 1998]. In liver and pancreatic tissues Tcf7l2 underpins glucose homeostasis [Norton et al., 2014], with intronic mutations that reduce expression of Tcf7l2 being associated with Type II diabetes [Grant et al., 2006]. One mechanism by which Tcf7l2 achieves cell-type specific effects is by binding at different genomic locations, hence regulating a different set of target genes. Across 6 human cell lines (5 endodermal, 1 epithelial) Tcf7l2 bound a largely disparate set of sites, with only 1,800 out of 116,000 total Tcf7l2 binding sites shared between the 6 [Frietze et al., 2012]. Supporting the idea of a grammar, different cofactor motifs tend to be enriched at cell-type specific binding sites. Similarly, in [Frietze et al., 2012] the Foxa2 and Hnf4*α* motifs are enriched in a hepatocyte cell line (Hnf4*α* appears to function with Tcfs in hepatocytes [Norton et al., 2014]), while in an adenocarcinoma cell line it appears that the Gata3 motif helps bind Tcf7l2 when its own motif is absent.

The non-random distribution of sequences in the genome, however, makes it difficult draw further inferences from features enriched at transcription factor binding sites. Two transcription factors could bind together because they control a similar set of genes, rather than because they stabilise each others binding. On the other hand, particularly strong arrangements of transcription factors could cause ectopic activation and be selected against, resulting in native enhancers being comprised of weaker than possible arrangements of motifs as they provide a sharper response to external signals [Farley et al., 2016]. As such the most informative arrangements of motifs for detecting interaction effects are likely under-represented in the genome. Furthermore, it is not clear how much of the binding of a transcription factor at a specific location is due to the sequence immediately surrounding it. Each site is an unique position in the genome, and could be influenced by the chromatin organisation in the region, looping and interactions with distal regions, and impact of transcription in the area. For example, ChIP-seq experiments performed on livers of mouse F1 crosses have detected the impacts of genomic variants up to 10kb away from binding sites [Wong et al., 2017]). Additionally, detecting binding sites is usually done through chromatin immunoprecipitation, which uses the same cross linking step as for detecting 3D interactions between distal genomic segments (looping). Without careful titration of this reaction one cannot be sure it is only direct transcription factor with DNA interactions, and not some larger complex, that one extracts [Teytelman et al., 2013]. This unavoidable combination of complex genomic features confounding measurement, removal of distal positional information, and biased sequence distribution, means that one is limited in the ability to determine transcription factor binding logic from computational analysis of genomic binding patterns.

To assess Tcf7l2 binding logic while avoiding the complexity of inferring from sites bound across the genome, we have developed an approach to measure Tcf7l2 binding to thousands of 99-bp variable sequences transplanted into fixed, defined genomic loci using a quantitative DamID assay [Vogel et al., 2007, Szczesnik et al., 2019]. This strategy allows us to detect differences in binding induced by minimal, designed sequence alterations while controlling for effects of the surrounding DNA sequence, thus enabling us to determine causal relationships between DNA sequence and Tcf7l2 binding. Importantly, our assay is performed in a native cellular chromatin context, allowing us to account for effects of chromatin organisation and interactions with other proteins that are missing in *in vitro* binding assays.

Using our novel approach, we find that while *in-vivo* Tcf7l2 binding is dependent on the presence and match of its *in-vitro* motif at individual binding sites, Tcf7l2 binding also varies dramatically based on the sequence surrounding it, and cell type-specific Tcf7l2 binding at genomic loci can be partially recapitulated by the local surrounding sequence. Particularly, the presence of Oct4 and Klf4 motifs favours Tcf7l2 binding in mESCs, and this effect is strongest when occuring within an adjacent 20 to 50-bp region and oscillates approximately every 10.8-bp shift in distance between the Tcf motif and cofactors. This effect is strongest surrounding when the Oct4 motif occurs as apart of the joint Sox2-Oct4 motif [Chen et al., 2008], and particularly helps promote binding in inaccessible chromatin which is otherwise refractory to Tcf7l2 binding. This novel high-throughput DamID assay provides a powerful platform to determine local DNA sequence grammars that causally influence transcription factor binding.

## 5 Results

We developed an assay for measuring Tcf7l2 binding to thousands of pre-determined DNA “phrases” at a specific genomic locus (Figure 1 A). Briefly, a library of synthetic oligos containing a 99-bp variable region (phrase) flanked by short constant sequences used as primers is integrated into specific genomic locations in mouse embryonic stem cells (mESCs) by CRISPR/Cas9 based homology-directed repair. Previous work has shown that 20-40% of alleles will have site-specific integration of one of the variable phrases [Hashimoto et al., 2016, Rajagopal et al., 2016]. Binding of Tcf7l2 to each integrated phrase is measured by DamID [Vogel et al., 2007] using doxycycline-inducible ectopic expression of a genomically integrated fusion protein of Tcf7l2 and the N126A mutant of Dam, which we have recently shown to allow accurate measurement of Tcf7l2 binding genome-wide with reduced off-target methylation as compared to the wild-type Dam enzyme [Szczesnik et al., 2019]. A Dam methylation site (GATC) is located directly adjacent to the site of phrase library integration. The genomic DNA is separately digested with restriction enzymes specifically recognising unmethylated GATC (DpnII) or methylated GATC (DpnI) and then PCR amplified with primers designed to only capture genomically-integrated, undigested phrases. These two pools of only methylated or unmethylated sequences are then sequenced by Illumina Next-generation sequencing (NGS), and the relative abundance of a particular phrase between the two pools is used to estimate the level of Dam methylation, and hence of Tcf7l2 binding, to the integrated phrase.

**Figure 1:**
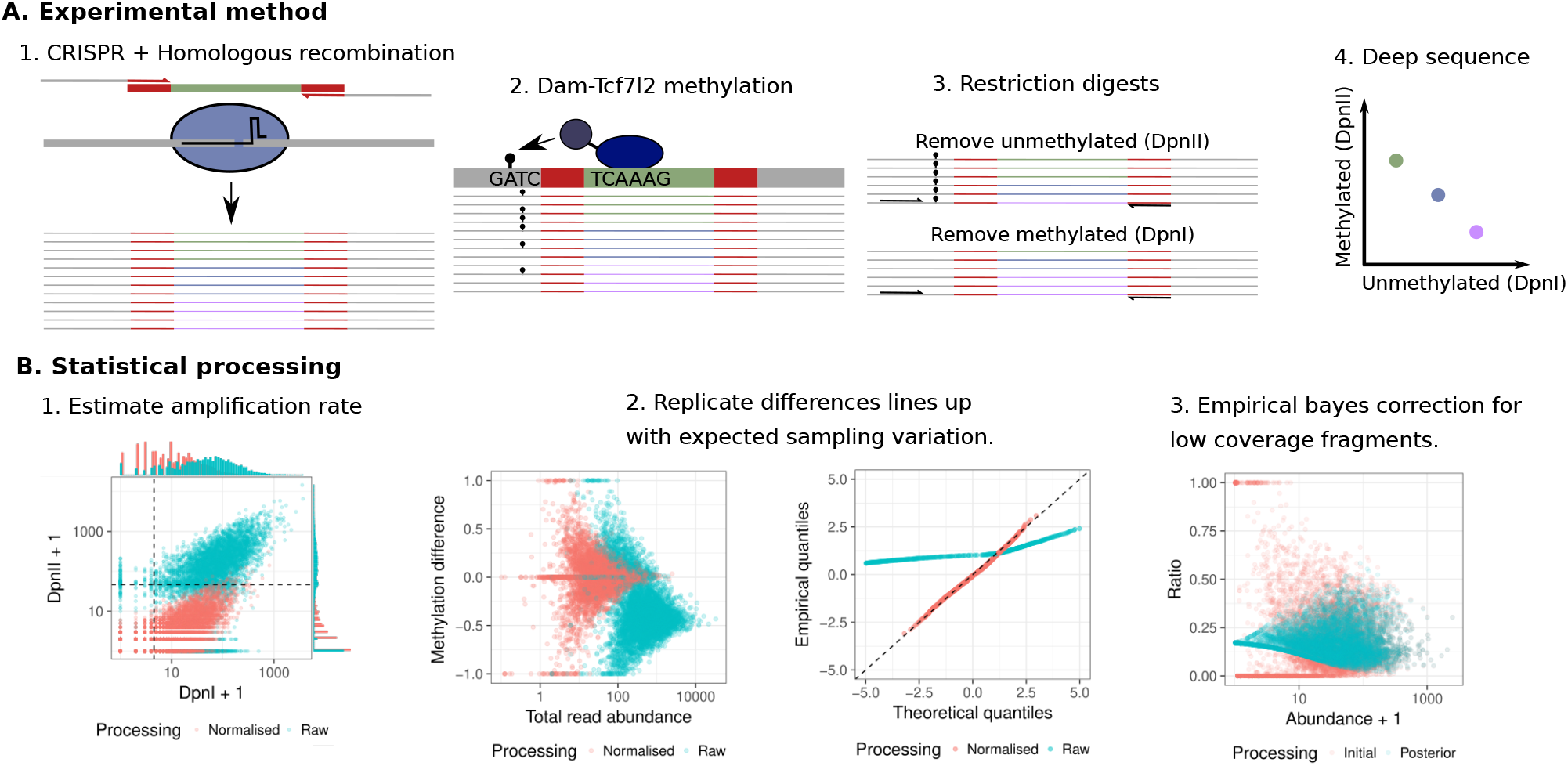
Methods used for assaying transcription factor binding to variations in a specific genomic locus. A) The phrase library is integrated using CRISPR/Cas9 and homologous recombination into a specific locus. Binding of Dam-Tcf7l2 results in methylation of a GATC adjacent to the site of integration. Following genomic DNA extraction, two pools of completely methylated or unmethylated are created with methylation specific restriction enzymes, which are then deep sequenced and mapped back to the initial library. The relative enrichment of a fragment in the two pools indicates the level of Tcf7l2 binding. B) A negative-binomial dropout model is fit to the data to estimate the degree of amplification in the resulting reads, and hence to estimate the number of initial methylation events that occured. Following normalisation, the distribution of variability between replicates closely follows the expected sampling variability. Finally, a beta-binomial empirical bayes model is fit to give a posterior distribution for each phrase’s methylation rate.

We initially designed and screened (by Tcf7l2 DamID) a library of 12,000 phrases. Half of this library was comprised of 99-bp genomic sequences that show variable Tcf binding. Cohorts of phrases were sampled from ChIP-seq peaks bound in either Tcf7l1 ChIP-seq in mESCs or Tcf7l2 ChIP-seq in intestinal endoderm cells, or in both (1200 phrases each; see Methods and Supplementary Figure 1). If present, any Tcf motif was included in the sampled 99-bp phrase, and we specifically sampled half the cohort from ChIP-seq peaks lacking any clear Tcf motif (half of each group). The final cohort of phrases contained unbound Tcf motifs (>10kb from any Tcf ChIP-seq peak) in regions of open chromatin near marks of active enhancers (H3K27ac ChIP-seq peaks) (2400 phrases). The remaining 6,000 phrases were generated *de novo* from different arrangements of binding motifs for a set of transcription factors we deemed likely to influence the binding of Tcf7l2, based on published protein-protein interactions or motif enrichment adjacent to Tcf binding sites (see Methods for details) [Frietze et al., 2012], [Norton et al., 2014], [Cole et al., 2008]. This library was first integrated in 2 biological replicates into the inert Rosa26 locus, which resides in natively accessible chromatin [Zambrowicz et al., 1997]. DamID on each of the replicates was done with both Dam-Tcf7l2 and unfused Dam, which has been shown vary with chromatin accessibility [Kladde, 1992] and otherwise provides a control for differences in Dam methylation rates between phrases.

We then developed a computational pipeline to normalize data arising from this experiment. Following alignment and counting we noticed that the read counts in the DpnI and DpnII digested samples did not follow the unimodal distribution expected of a large variable library, suggesting a a bottleck in unique methylation events followed by PCR duplication (Figure 1 B1). To estimate the number of initial genomically-integrated phrases captured by the sequencing we developed a zero-inflated negative binomial model (see Methods for details). Briefly, latent genomic counts are modelled as a negative binomial distribution, and observed sequences in each sample from a poisson distribution stemming from a linear amplification of these latent counts (see Supplementary Data for values). Following parameter estimation, the latent genomic counts closely follow the expected binomial distribution between replicates, indicating that the vast majority of difference between replicates stems from sampling variability (Figure 1 B2). We observed, however, that many phrases have few genomic counts (<100) which results in a highly variable estimate of the underlying methylation rate. To account for this variability without drawing a hard threshold that would discard useful information, we use a beta-binomial empirical bayes model to calculate the variance of each phrase’s methylation rate, utilising the distribution of methylation rates across the whole library to bias estimates of extreme values with low read count towards the population mean. Beta-binomial models are used to the same effect in reducing the false positive rate of detecting methylated cytosines with few supporting reads in genome-wide bisulfite sequencing [Sun et al., 2014]. This model yields a posterior distribution of methylation rate for each fragment, which we use for further inference (Figure 1 B3). In practise this adds a few pseudocounts (1 to 14, see Supplementary Data for values) to every sample, biasing low coverage fragments towards the overall population mean but not affecting the high coverage fragments. Overall, this computational pipeline allows us to accurately quantify the uncertainty in estimating Tcf7l2 binding for each phrase of a 12,000 phrase library integrated into a fixed genomic context.

We next proceeded to assess what features govern Tcf7l2 binding, by comparing to the methylation rate seen with the unfused Dam and extracting significant features through logistic regression with cross-validation (glmnet, [Friedman et al., 2010]). For the 6,000 phrases derived from native genomic sequences, specificity for cell-type specific binding of Tcf7l2 is retained when these sequences are transplanted to an open chromatin site in mESCs, with phrases bound in mESCs bound more strongly than those only in bound in intestinal endoderm or completely unbound (0.23 vs 0.046 effect sizes, unbound phrases are at 0; Figure 2 A). Presence of a Tcf motif is a strong predictor of Tcf7l2 binding in our assay, as phrases originating from Tcf ChIP-seq peaks but lacking a Tcf motif tend to show weak or absent Tcf7l2 binding, as compared with Tcf motif-containing sites within the same ChIP-seq binding profile (ESC 0.36, IE 0.27, Both 0.27 effect sizes). In particular, out of the phrases only bound in intestinal endoderm, only those with a Tcf motif tend to allow for binding when transplanted into the Rosa26 locus (0.27 vs 0.046 effect sizes). Within the set of phrases derived from sites unbound by Tcf7l2 in mESCs, the presence of enhancer marks in mESCs is a strong predictor that they will not acquire binding in the Rosa26 locus, irrespective of whether they exhibit Tcf7l2 binding in intestinal endoderm or not (IE −0.24 effect size, unbound at 0). Overall, the preferential binding of Tcf7l2 to Tcf motif-containing phrases derived from sites with native mESC Tcf7l2 binding over those with Tcf motifs but without native mESC binding lead us to conclude that the sequence immediately surrounding a Tcf motif is strongly responsible for regulating the binding of Tcf7l2 *in vivo*.

**Figure 2:**
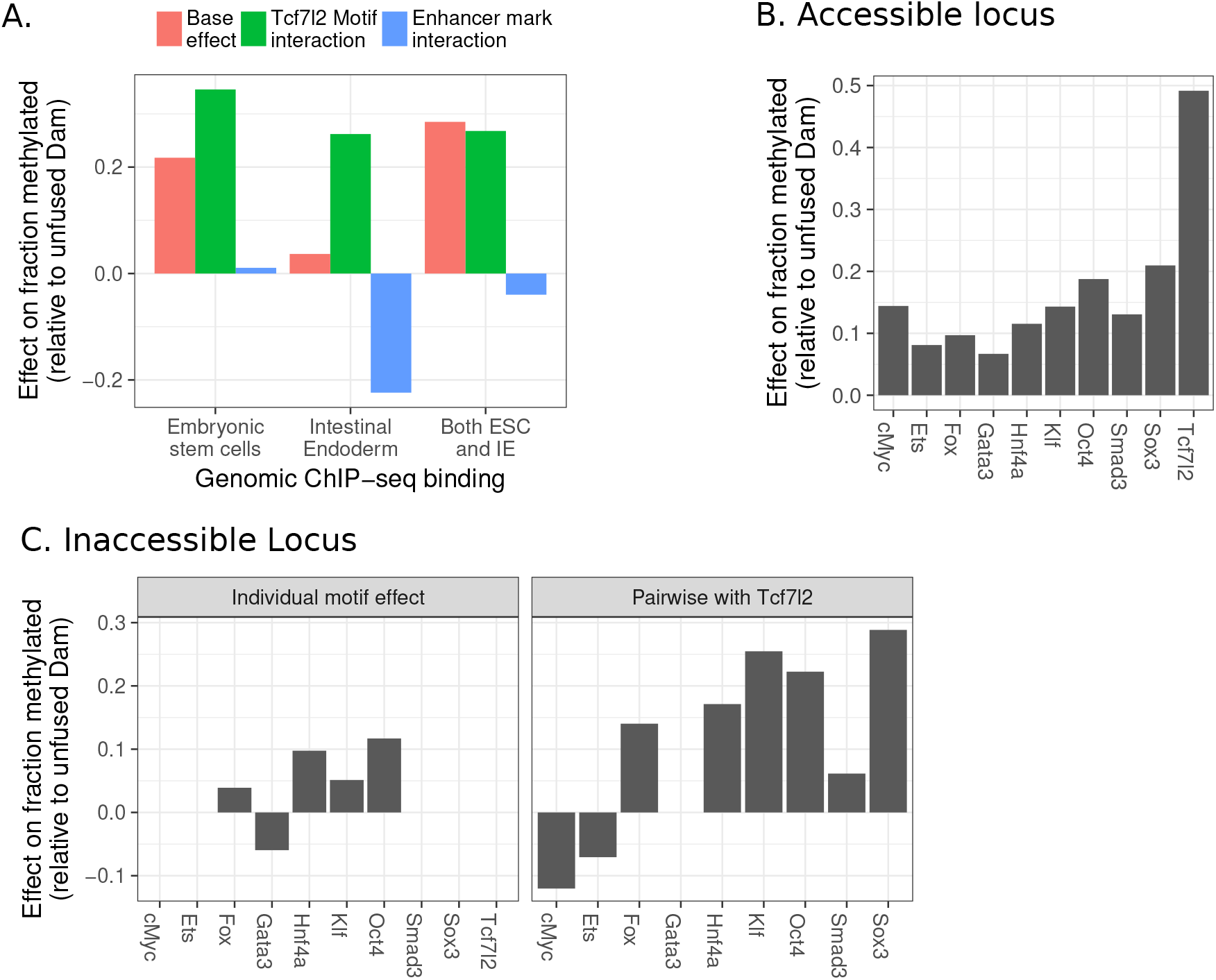
99-bp of local sequence regulates binding of Tcf7l2 to its motif across different genomic sites. A) Logistic regression effect size of genomic features on phrases transplanted to the accessible locus Rosa26. B) Logistic regression effect size for the presence of cofactor motifs after integration of the sequence library in the accessible locus. C) Logistic regression effect size for the presence of cofactor motifs, either individually or as pairwise interaction with the presence of the Tcf motif, after integration of the phrase library in the inaccessible locus uCD8. This specific sample was measured with wild-type Dam, instead of the N126A variant used elsewhere.

For the 6,000 phrases containing different motif arrangements, we calculated the effect of the presence of each motif on Tcf7l2 binding. We found that the presence of the Tcf motif had the strongest effect on Tcf7l2 binding (0.49 logistic regression effect size), and other putative cofactors had weaker but still positive effects on Tcf7l2 binding (~0.1 to 0.2 effect size; Figure 2 B). To distinguish effects on Tcf7l2 binding driven by protein-protein interaction as compared to those driven indirectly by induction of chromatin accessibility adjacent to the Tcf motif, we integrated this 12,000-phrase library into a genomic locus with minimal native chromatin accessibility in mESCs (upstream of the T-cell-specific CD8 gene: uCD8). Due to the low signal detected with Dam-N126A for this library in this locus, we used the wild-type version of the Dam enzyme which gives stronger signal, however at the cost of being strongly confounded by changes in chromatin accessibility. We found that, in this natively inaccessible locus, Tcf7l2 binding required the presence of specific cofactors, particularly Oct4, Klf4, or Sox2 (0.25, 0.22, 0.30 effect sizel; Figure 2 C). Thus, in this controlled assay, Tcf7l2 binding is impacted by adjacent motifs, and these motifs become more important when phrases are integrated into a locus without surrounding chromatin accessibility.

A deeper analysis of these interactions would require single phrase-resolution data of this library, which we were unable to obtain due to insufficient data coverage which is limited by the incomplete efficiency of CRISPR/Cas9-based homology-directed repair and feasible cell culture volumes. Out of the 12,000 phrases, we observed integration of only 3,000-4,000, and only a few hundred (9%) had sufficient coverage for accurate estimates of their true methylation rate (Figure 3 A), limiting analysis to population trends and preventing the detection of sparser levels of binding in the inaccessible locus. In summary, the results of this 12,000 phrase library reveal that the sequence surrounding a Tcf motif imparts a degree of cell type-specificity to Tcf7l2 binding. This is partially dependent on the presence on cofactor motifs; however technical limitations prevented us from examining spatial dependencies and interactions with the local chromatin environment that underlie these cofactor interactions.

**Figure 3:**
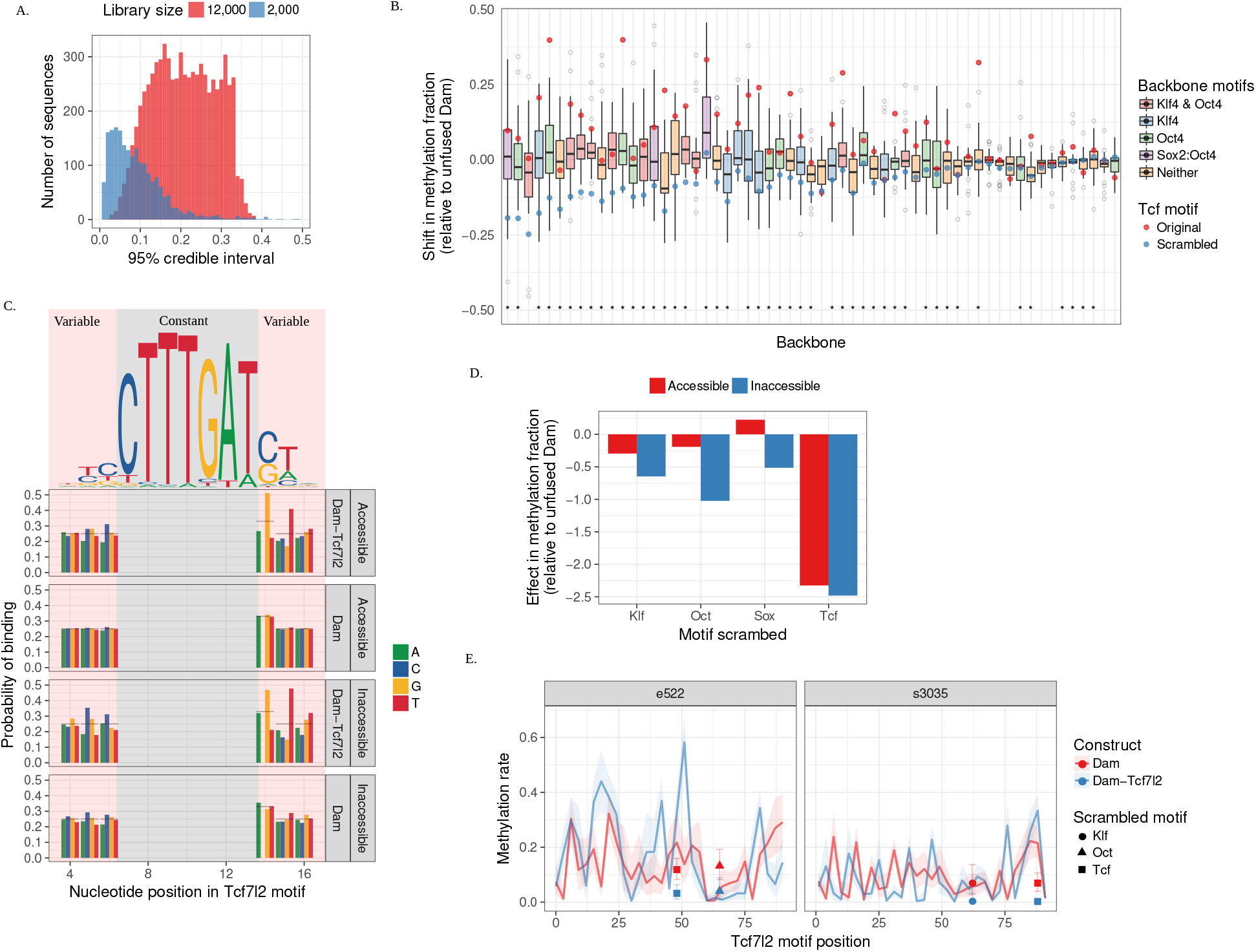
Individual phrase level resolution of Dam-Tcf7l2 binding shows influence of motif strength and cofactor motif presence. A) Comparison of 95% posterior credible interval of fraction methylated for the 12,000 and 2,000 phrase libraries with Dam-Tcf7l2 in the accessiblel locus. B) Spread of Tcf7l2 binding (Dam-Tcf7l2 relative to unfused Dam) as the Tcf motif is rolled across each backbone. Scrambling the Tcf motif (red dot) substantially decreases binding relative to original (* indicates p < 0.05). C) The two 5’ and 3’ nucleotides flanking the Tcf motif consistently explain some of a variability in binding, and are consistent with lower informative bases in the estimated motif for Tcf7l2 from protein binding microarrays. D) Logistic regression effect of scrambling each motif across the whole library. For raw values see Supplementary Figure 2. E) Example of footprint for the rolling Tcf motif disrupting the Oct4 (e522) or Klf4 motif (s3035) to the same level as scrambling the motif. Shaded region / error bars show 95% posterior credible interval.

To investigate the adjacent motifs and spatial determinants governing Tcf7l2 binding at higher resolution, we designed a 2,000 phrase library that systematically varied the position of the Tcf motif across a set of 59 backbone sequences. By reducing the number of phrases, we posited that we would increase coverage of each phrase and thus enable phrase-resolution analysis. Backbones were chosen from both the native genomic and synthetically generated phrases in the initial library so as to span a range of Dam-Tcf7l2 methylation rates across both accessible and inaccessible chromatin loci (see Methods for details). For each backbone, phrases were designed with a scrambled version of the initial Tcf motif, and a set of phrases was designed in which the most informative, core part of the Tcf motif (CTTTGAT) was “rolled” across each backbone in 3bp increments, replacing the sequence that had been in that position. We also included phrases for each backbone in which we scrambled motifs for Oct4, Klf4, and Sox2 which were identified from the initial library as likely influencing Tcf7l2 binding (see Methods for scrambled sequences). We integrated this 2,000-phrase library into the Rosa26 and uCD8 genomic loci in two biological replicates in mESCs, and performed DamID with Dam-Tcf7l2 and unfused Dam as for the previous library, with the difference that a methylation site (GATC) was included on both sides of the integrated phrase in order to reduce possible spatial effects from distance of Dam-Tcf7l2 to the GATC.

Integrating this smaller phrase library vastly reduced the uncertainty in estimated methylation rate for individual fragments (Figure 3 A), providing much higher resolution data (9% to 65% increase in the number of phrases with a 95% credible interval less than 0.1; >99% phrases were recovered). This was due to sampling many more unique genomic instances of each phrase, and since the variability between replicates remained consistent with the beta-binomial model (at a 0.05 cutoff, 0.065 of the methylation rates of the second replicate fell within the posterior distribution of the first replicate) the concordance between replicates increased proportionally. Thus, we were able to perform individual phrase-level analysis of Tcf7l2 binding for the majority of the 2,000 phrases.

We first looked at Tcf7l2 binding in the accessible locus. Consistent with our findings in the initial library, scrambling the Tcf motif led to significantly decreased Tcf7l2 binding in backbones (46/59 at p < 0.05; Figure 3 B). When comparing Tcf7l2 binding in phrases from the same backbone, we found substantial variation as the Tcf motif is rolled across each backbone (standard deviation range 0.029 - 0.16; mean of 0.084; Figure 3 B). This intra-backbone variation is significantly larger than the standard deviation in the estimate of each phrase’s methylation rate (only 1.5% of phrases had a higher rate), indicating that the assay is sensitive enough to detect changes in binding resulting from rolling the Tcf motif. The few backbones that did not have a significant drop in methylation upon scrambling the Tcf motif also had a lower variability in Tcf7l2 binding as the Tcf motif was rolled (mean standard deviation of 0.06 vs 0.09). In fact, on average, Tcf7l2 binding in the phrase with the most unfavourable Tcf motif position within a backbone was equivalently low to the phrase with scrambled Tcf motif, indicating that the sequence context is a strong determinant of binding (22/59 backbones had significantly lower p < 0.05 methylation at the worst position, and in no instance was the scrambled Tcf motif significantly lower than the worst position; Figure 3 B). We conclude that the location of a core Tcf motif relative to surrounding local sequence plays an important role in determining Tcf7l2 binding strength. We thus turned to investigating patterns in these Tcf motif rolling experiments to determine the key features underlying Tcf7l2 binding logic.

We find that one major cause of the variability in Tcf7l2 binding as the motif is rolled across each backbone is the change in nucleotides flanking the core Tcf motif. While we rolled the 7 nucleotide core Tcf motif, the position weight matrix that best explains Tcf7l2 in protein binding experiments [Badis et al., 2009] contains contributions from two nucleotides on either side of the core motif. By calculating the average Tcf7l2 binding across all possible base identities in the flanking positions, we find that Tcf7l2 binding strength in our assay correlates with the optimal nucleotide identities of full Tcf7l2 position weight matrix – a 3’ guanine followed by thymine, and a slight 5’ cytosine or thymine preference (Figure 3 C). We note that 3’ cytosines are not observed as they were replaced with G to avoid the creation of an extra GATC methylation site. A logistic model of the contribution of flanking nucleotide position to Tcf7l2 binding found that including the effects of di-or tri-nucleotides did not improve the model fit (increases binomial deviance from 0.082 to 0.086), indicating that independent contributions of single nucleotides are sufficient to explain variation in binding strength from flanking nucleotides. This finding rules out effects of new motifs being reproducibly formed from rolling the Tcf motif, as they would result in a model that prefers base-pair dependencies and would likely differ from the in-vitro binding preference. Tcf7l2 binding differences resulting from these differences in motif affinity explain 35 - 40% of the variability in binding within each backbone (see later modelling in Figure 4 A). Motif affinity does not, however, predict any differences in binding affinity between backbones, since the flanking nucleotides are uncorrelated with other sequence features.

**Figure 4:**
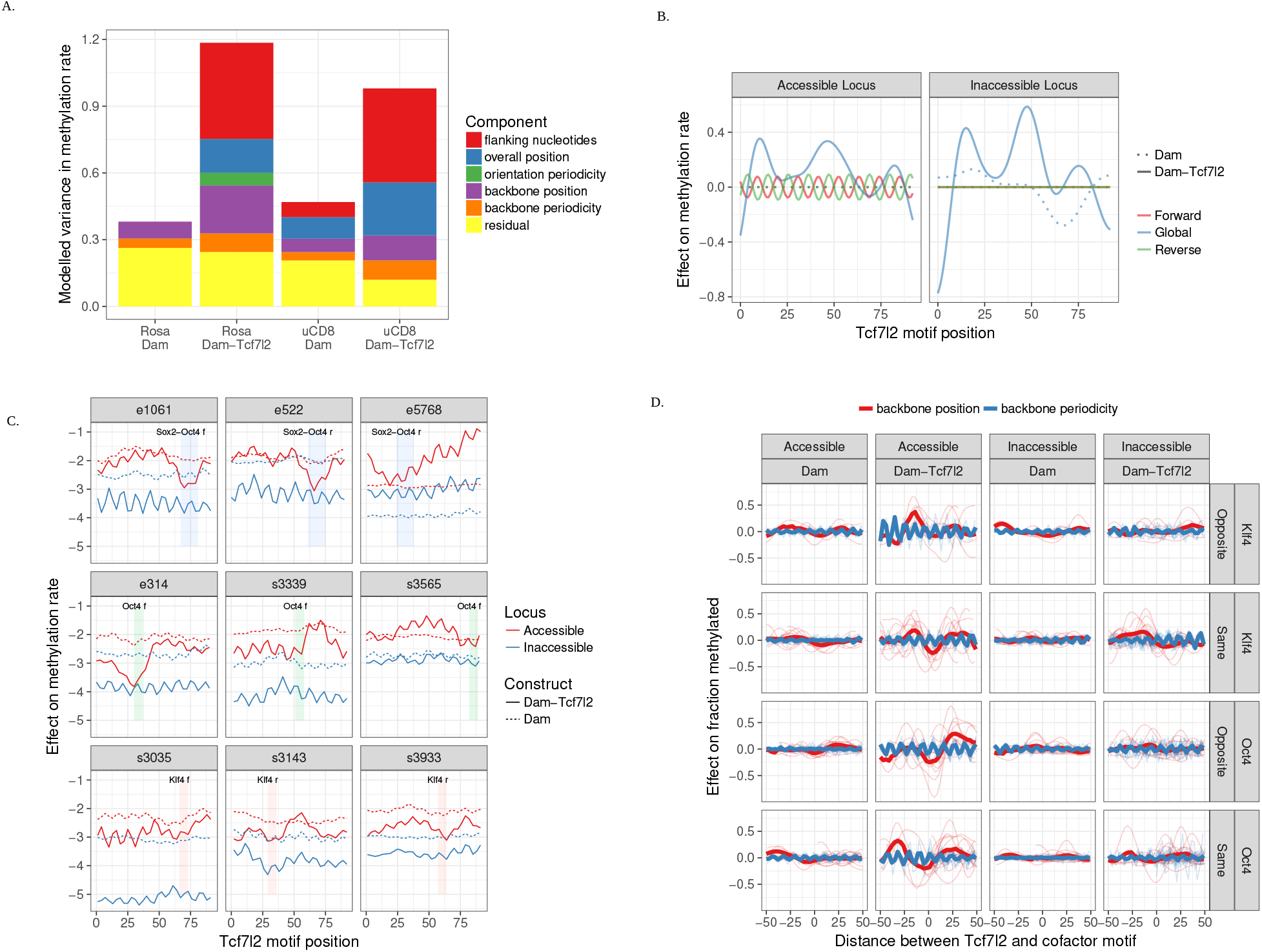
Tcf7l2 binding depends on locus and cofactor dependent spatial interactions. A) Intra-backbone variance in methylation rate explained by gaussian process components across both accessible and inaccessible loci, and Dam-Tcf7l2 and unfused Dam. B) Estimated effect of the backbone-invariant smooth and orientation specific periodic effects. C) Observed methylation components extracted from the gaussian process for a representative set of backbone sequences. Shown is the sum of the backbone specific components: constant, smooth, and periodic. D) Aggregate profiles of the backbone specifc smooth and periodic components for Dam-Tcf7l2 or unfused Dam methylation, group by cofactors present. Phrases are centered at midpoint of the respective cofactor motif, and calculated to the midpoint of the Tcf motif for each of the possible orientations. Sox2-Oct4 joint motifs are excluded as they would exaggerate the average signal compared to other Oct4 sites.

Much of the remaining variation in positional Tcf7l2 binding appears dependent on the presence of neighbouring motifs, as backbones with the highest binding rate contained either a Klf4 or Oct4 motif (Figure 3 B), and shuffling these motifs also tended to reduce binding (Figure 3 D). Plotting the Tcf7l2 binding strength as the Tcf motif rolls along a backbone, we identify striking patterns of reduced Tcf7l2 binding resembling “footprints” coinciding with phrases in which the Tcf motif disrupts an underlying Oct4 or Klf4 motif, to the same degree as scrambling these motifs (Figure 3 E). Since the Tcf motif is rolled by 3bp, any motif longer than 3bp should be detected. Nonetheless, we observed no instances of loss of Tcf7l2 binding for several adjacent positions occuring for any other known motifs, although we cannot rule out that other motifs would show this effect if we had rolled the Tcf motif across a larger cohort of backbones. Even within the set of backbones containing Oct4 or Klf4 motifs, we observed substantial backbone-specific variation in the magnitude of Tcf7l2 binding and the loss of such binding upon disruption or scrambling of these cofactor motifs (Supplementary Figure 2). The strongest effects were localised around backbones containing an Oct4 motif as part of a joint Sox2-Oct4 motif, which hints that much of this variability stems from differences in binding affinities of these cofactors between backbones. However, because of the low numbers of instances of either Oct4 or Klf4 motifs in the 59 backbones and the fact that we did not vary the strengths of these motifs in a controlled way, we cannot make strong conclusions about the role of cofactor motif strength.

Having identified significant roles of the extended Tcf motif and the presence of adjacent Oct4 and Klf4 motifs in modulating Tcf7l2 binding, we sought to build a computational model to explain Tcf7l2 binding as a function of its primary motif and position within the backbone. In order to extract such positional effects we use gaussian process regression (see Methods for details). This technique allows us to fit classes of non-linear functions based on the changing position of the Tcf motif. Importantly, such non-linear functions allow the model to generalise between different Tcf motif positions without treating each position as independent and without pooling across several positions. As a result, gaussian process regression should allow us to more easily identify reproducible co-factor interactions that lack fixed spacing.

The overall model attempts to explain Tcf7l2 binding across the 2,000 phrases by identifying an optimal linear combination of non-linear spatial effects of Tcf motif position, flanking nucleotides, average methylation per backbone, and each sequence’s unique methylation (i.e. unexplained residuals). The uncertainty in estimated methylation rate for each phrase is propagated to this model through a logit-normal approximation to the posterior beta distribution. The non-linear spatial effects of Tcf motif position are separated into those that are backbone-invariant, either across all backbones or those sharing the same Tcf motif orientation, or spatial effects unique to each backbone. These spatial effects are captured by two different classes of functions. The first class of functions are designed to identify stretches of higher or lower Tcf7l2 binding, and are modelled by a radial basis kernel, which fits smoothly varying functions and is parametrised by a length scale that controls how quickly the function varies. The second class of functions are similar in nature but oscillate periodically, and hence are additionally parametrised by a periodicity. The hyperparameters of the kernels and the linear weights for the relative contribution of each component are found by optimising the marginal likelihood through gradient descent (periodicity is first estimated by a search over all integers prior to gradient descent).

Using this gaussian process regression technique, we find that the model that best fits the observed Tcf7l2 DamID data in both accessible and inaccessible chromatin loci contains inputs from multiple distinct components (Figure 4 A). The most salient individual component influencing intra-backbone variation in Tcf7l2 binding (i.e. after excluding averaged differences in Tcf7l2 binding across backbones) is Tcf motif strength as determined by the identity of the four nucleotides adjacent to the Tcf7l2 core motif which was rolled, which explains 35 and 40% of Dam-Tcf7l2 methylation variability in accessible and inaccessible loci respectively. The various spatial effects of Tcf motif position explain a further 45% of the variation in binding across both loci. Within these, the backbone-invariant spatial effects which depend on overall position and orientation of the Tcf motif across all backbones explain 20 and 25% of variable methylation in accessible and inaccessible loci. Backbone specific spatial effects, those that only generalise across Tcf motif positions in the same backbone, explain a further 25 and 20% of variable methylation in accessible and inaccessible loci. After accounting for Tcf motif strength, spatial effects stemming from motif position, and uncertainty in the estimated methylation rate, there is a residual per phrase binding variability of 20 and 15% in accessible and inaccessible loci. Thus, a gaussian process regression model with a minimal set of features is capable of explaining the majority of variation in Tcf7l2 binding across the 2,000 phrases.

The variability in Dam-Tcf7l2 methylation is considerably higher than that of unfused Dam seen when rolling the Tcf motif (4X higher in the accessible locus, 2X higher in the inaccessible locus, Figure 4 A). Unfused Dam methylation has much higher unexplained residuals (accessible 75%, inaccessible 50%), with smaller contributions from backbone-specific effects (accessible 25%, inaccessible 20%) and an effect of overall position of the Tcf motif only in the inaccessible locus (15% of variability). The inaccessible locus also has ~10% of variance explained by contribution of flanking nucleotides, however in this case it does not recapitulate the *in-vitro* Tcf position weight matrix, and instead appears to resemble a putative weak Sox2-Oct4 motif: 5’ Cs and 3’ A-T-G bias (Figure 3 C). These results suggest that the unfused Dam signal in inaccessible chromatin is capturing subtle changes in chromatin accessibility arising from the creation of weak alternative motifs while rolling the Tcf motif – the Tcf, Sox2, and Oct4 motifs all share a core TTTG stretch, making it difficult to definitively assign the most likely binding factors to sites with weak position matrix weight matches. The effect of cofactors influencing accessbility can also be seen in the smooth backbone-specific effect with unfused Dam, which while weaker is correlated with Dam-Tcf7l2 (accessible: 0.35, inaccessbile: 0.18; Supplementary Figure 3). In sum, the gaussian process regression model is less effective at determining the causes of variation in unfused Dam methylation, likely because the input features have been tailored to predicting Tcf7l2 binding variation. This finding reinforces that the model is learning features specific to Tcf7l2 binding and not to confounders introduced by the DamID method.

Unexpectedly, the overall position of the Tcf motif in the locus has an effect on Tcf7l2 binding. The spatial effect of the Tcf motif position across all backbones (irrespective of orientation) shows a similar pattern in both accessible and inaccessible loci: the length scale optimises to ~8-bp, and extracting the estimated function (Figure 4 B) shows that Dam-Tcf7l2 methylation is highest when the motif is located within the middle of a backbone or towards either end with dips in intervening regions. Since this effect is so similar for Dam-Tcf7l2 methylation across both loci, we posit that a likely cause is steric constraints on how well Dam-Tcf7l2 can methylate the GATC sites located adjacent to the ends of each integrated phrase, rather than changes in Tcf7l2 binding. Unfused Dam in the inaccessible locus also shows an effect of the overall Tcf motif position (Figure 4 B), however this only mimics an overall downward trend of Dam-Tcf7l2 and not the middle and end peaks, presumably because unfused Dam is not binding to the Tcf7l2 motif. Thus, we do identify one feature that is best explained as an artifact of the DamID method – Tcf7l2 is on average more adept at methylating GATC’s with particular distance constraints.

When backbone-invariant effects are calculated only across backbones that share the same Tcf motif orientation, the model identifies a periodic function only in the accessible locus (Figure 4 B). The periodicity parameter optimises to every 10.8 bases, which is close to the estimate of a DNA helix rotation (10.4 - 10.6) [Wang, 1979, Rhodes and Klug, 1980, Klug and Lutter, 1981]. Interestingly, the periodic component for the two Tcf motif orientations are completely out of phase. Since Tcf7l2 binding introduces a large bend in the DNA (90 - 127 degrees) [Love et al., 1995, Giese et al., 1995], a possible explanation is that DNA bending in the Rosa26 accessible locus is more energetically favourable in one direction. This would favour Tcf7l2 binding sites that are in-phase with one another with respect to the rotation of a DNA helix, since these would all bend in the same direction.

Lastly, we investigated the backbone-specific spatial effects. The optimal model includes input from both smoothly varying and periodic backbone-specific components with similar hyperparameters as the global and orientation effects (~8 rbf length scale and 10.8 periodicity), in both the accessible and inaccessible loci. The smoothly varying, periodic, and constant backbone-specific effects are extracted and summed up for a set of representative backbones (Figure 4 C).

The smooth component captures the footprints of reduced binding as an underlying cofactor motif (Oct4 or Klf4) is disrupted (compare backbones e522 and s3035 in Figure 4 C with Figure 3 E). Additionally, the smooth components captures regions of higher Tcf7l2 binding near the cofactor motifs the Oct4 and Klf4 motifs (2nd and 3rd row in Figure 4 C), which tend to spread over an ~20 to 50-bp region adject to the cofactor motifs. It is possible that another motif is being disrupted after these stretches of higher binding, however since we could not reliably identify any other motifs at these locations we believe that it represents the effect of Oct4 or Klf4 promoting adjacent Tcf7l2 binding within this window.

The backbone-specific periodicity captures a reproducible oscillation in binding strength estimated to occur every 10.8-bp. We hypothesize that backbone-specific periodicity in Tcf7l2 binding arises from interaction between Tcf7l2 and cofactors such as Oct4 that is strengthened when both factors reside on the same side of the DNA helix. The estimated periodic effects are similar for Dam-tcf7l2 across both loci (0.35 correlation, vs 0.075 - 0.14 to unfused Dam; Supplementary Figure 3), indicating that it is detecting oscillating patterns specifically promoting adjacent Tcf7l2 binding within this window.

The backbone-specific smooth and periodic effects across different backbones tend to align when aggregated across different backbones based on the relative position between the Tcf and the Oct4/Klf4 cofactor motif (Figure 4 D). There is a large variability in the strength of these backbone-specific effects across different backbones – the three backbones that contain a joint Sox2-Oct4 motif possess the strongest such effects (first row of Figure 4 C). Since there are only 3 such backbones, they are excluded from the aggregate profiles in Figure 4 D to avoid exaggerating them, but the estimated positional effects overlap for the shared part of the orientation and gap spacing (Supplementary Figure 5) These three Sox2-Oct4 motif containing backbones which show strong dips when the Sox2-Oct4 motif is disrupted, longer than average stretches of improved Tcf7l2 binding nearby, and strong backbone-specific periodic effects, with maximal Tcf7l2 binding when Tcf7l2 is separated from Oct4 by 27-29, 37-39, 48-50, and 61bp (relative to the midpoint of the Tcf7l2 and Oct4 motifs, and similar across all four orientations of motifs). Thus, aligning backbones by the distance between the Tcf7l2 motif and Oct4/Klf4 co-factor motif reveals that the smooth and oscillating effects are consistent, and are accentuated in the three Sox2-Oct4 motif-containing backbones where co-factor binding strength is likely to be strongest.

In summary, in-depth analysis of this collection of 2,000 phrases that examine Tcf7l2 binding logic at 59 Tcf motif-containing backbones reveals that Tcf7l2 binding is dependent on the binding of Oct4 and Klf4 motifs in a spatially dependent manner. Strongest Tcf7l2 binding occurs when the motifs are separated by 20–50 bp with oscillatory strength that matches the turn of a DNA helix. The strength of these effects appears dependent on the strength of the cofactor motif: it is strongest at joint Sox2-Oct4 sites, moderate at most other Oct4 and Klf4 sites, and weak at a few Oct4 and Klf4 motifs. Since Tcf7l2 binding is strongly dependent on sequence context, it is likely that similar effects exist that govern the binding strength of Oct4 and Klf4. Deconvolving all of these variables will require a substantially larger set of backbones that systematically vary in Oct4/Klf4 motif strength as well as relative Tcf7l2 position. The lack of other observed motifs in any observed footprints nor in the flanking nucleotides, suggests there are unlikely to be many other motifs that substantively influence mESC Tcf7l2 binding in the backbones analysed.

## 6 Discussion

The binding motif for Tcf7l2 has been well characterised *in-vitro*, however it is a poor predictor of the *in-vivo* binding of Tcf7l2 and lacks the ability to explain differences in cell-type specific binding [Frietze et al., 2012]. Here we use a combination of site-specific integration and *in-vivo* transcription factor binding measurement to show that a large contribution to the specificity of Tcf7l2 binding in mESCs is contained within the 99-bp of sequence surrounding a genomic Tcf motif, in particular the presence of Oct4 and Klf4 motifs. We also show that binding is strongly determined by the local chromatin accessibility, and that ectopic gain of Tcf7l2 binding occurs with sites usually only bound by Tcf7l2 in intestinal endoderm but located in inaccessible chromatin in mESCs. Thus, our results provide strong supporting evidence with an earlier classification of Tcf7l2 as a “migrant” transcription factor that is dependent on both local chromatin accessibility and on interactions with co-factors for binding [Sherwood et al., 2014].

A similar helical-dependent effect as shown here for Tcf7l2 with Oct4 and Klf4 has been observed previously in the *in-vitro* formation of a Lef1 (part of the Tcf family), Ets1, and CREB complex, which found that Lef1 promoted interactions between the flanking motifs in a way dependent on the phase of CREB in the DNA helix [Giese et al., 1995]. Our work extends this to show that DNA helix influenced co-factor binding can occur over several subsequent turns, and that such subtle effects of spatial positioning between transcription factor binding motifs play important roles in determining binding in a genomic context.

While the DNA binding domains of transcription factors are generally small and well folded, the remaining domains responsible for interactions with other proteins are often disordered or connected by flexible segments [Liu et al., 2006]. This flexibility should allow interacting domains of two adjacently bound transcription factors to interact across a range of spacings. This predicted flexibility in interaction distance is consistent with results from small scale enhancer reporter assays, [Erceg et al., 2014] which found that varying the relative spacing between the pMAD and Tin motifs can affect expression of a reporter gene integrated in a developing fruit fly (Drosophila) embryo. In one tissue, shifting from a 2bp to 8bp gap is enough to abrogate reporter expression, while in a different tissue it only halves expression. Importantly, in both cases this change is gradual: gaps of 4bp and 6bp exhibit intermediate function. Similarly, [Farley et al., 2016] tested variants of an enhancer that is active during sea squirt development (*Ciona intestinalis*) in an oligonucleotide reporter assay and found that small shifts in the distance between Ets and Zicl binding sites within it can change the strength of the enhancer. Our work extends upon these studies by showing that these spatially flexible co-factor effects span over a larger spatial region at least up to 50-bp and can influence binding of transcription factors.

Our finding that TF-TF interactions can occur over a range of distances but also change in strength in oscillating fashion by shifting spacing by 3-bp has implications for current approaches that model and predict TF binding. Typically, computational models of motif interactions are designed to allow for a constant effect irrespective of their relative positions, or to occur at an invariant spacing. For example, position weight matrices, linear regression, and support vector machines (i.e. [Ghandi et al., 2014]), all utilise sequence representations based on short subsequences or fixed gap lengths, that do not readily generalise across different gap lengths. Pooling across several different positions, for example as is utilised in convolutional neural networks for protein-DNA binding [Alipanahi et al., 2015, Kelley et al., 2016], similarly blurs over the smooth and oscillating spatial effects observed here. There are certainly cases in which TFs may require fixed spacing as with Sox2-Oct4 binding to a directly adjacent dual motif [Chen et al., 2008, Jolma et al., 2015]. It is likely, however, that cases like the Tcf7l2-Oct4 interaction we identify with oscillating and variable spacing are at least as typical. The covariance functions used here could be used directly as the kernel in a support vector machine to model such effects, while convolutional neural networks could utilise pooling layers across every Nth (e.g. every 10 or 11-bp) position rather than locally.

From a data standpoint, evaluating the effect of TF interactions across spacings requires many more data points than treating the interactions as spacing-agnostic or spacing-invariant. It is highly unlikely that we could have identified the Tcf7l2-Oct4 interaction patterns we found using genome-wide TF binding data. The genome does not have enough examples of these two motifs at varying spacings, and each genomic site has many other adjacent binding sites that would complicate modeling. Even with our approach using thousands of designed phrases, we are limited in power for building up an accurate model of Tcf7l2 binding. In several cases the residual methylation signal often appears to line up with the periodic components over a few positions, indicating that a more complex cooperative interaction exists within the data than is captured by a linear combination of spatial effects on Tcf7l2 binding.

Similarly, we lack enough observations to account for changing strength of Oct4 and Klf4 motifs (unlike for Tcf7l2 where we can accurately capture the strength of the motif through changes in flanking nucleotides). Particularly, the effect of Oct4 on Tcf7l2 binding in different backbones shows a range of magnitudes. This paucity of backbone variation may explain why the Sox2 motif significantly modulates Tcf7l2 binding in the larger first screen but not in the second screen with limited backbone examples. Klf4 motifs also showed variable strength in influencing Tcf7l2 binding across the second library. Additionally, some Sp1-like motifs – similar to the Klf4 motif – appeared like they might influence binding, but these effects were neither strong enough to produce a reliable “footprint” for specific instances, nor consistent enough to detect a consensus motif. A more complete and predictive understanding of Tcf7l2 binding logic would require rolling or varying the strength of Oct4 or Klf4 motifs across Tcf motif containing backbones as well as directly measuring binding of both Oct4/Klf4 and Tcf7l2 in the same set of backbones.

Our experimental design is also confounded by using ectopic expression of Tcf7l2 that is fused to an enzyme, so the altered levels of Tcf7l2 expression or altered function of the fusion protein may not perfectly mimic native Tcf7l2 binding. Compared to alternatives, such as the lossy ChIP assay, the gain in resolution offered by DamID makes this trade-off worthwhile when looking at individual loci, which occur at most twice in each cell.

Overall, this study demonstrates the power of massively parallel integration of DNA sequence variants into a controlled locus to address aspects of transcription factor binding logic that are difficult to address using observational genomic approaches such as ChIP-seq or in vitro approaches such as protein binding matrix arrays. In the future, this approach could be expanded to address co-binding logic by profiling binding of multiple transcription factors to the same collection of sequences, dynamic transcription factor binding by profiling binding in different cell-types, or combined with gene expression readouts to link transcription factor binding patterns to gene regulatory activity.

## 7 Author Contributions

T.S., J.W.K.H, and R.S. designed the experiments. T.S., L.C., and R.S. collected the data. T.S. analysed the data and wrote the code. T.S. and R.S wrote the manuscript.

## 8 Acknowledgements

This work is supported in part by a Human Frontier Science Program Young Investigator Grant (RGC0084/2014), the National Health and Medical Research Council of Australia (1105271) and the National Heart Foundation of Australia (100848).

## 9 Figures

## 10 Supplementary Figures

## 11 Methods

### 11.1 Lead Contact and Materials Availability

Further information and requests for resources should be directed to the Lead Contact, Dr Rich Sherwood

**Supplementary Figure 1:**
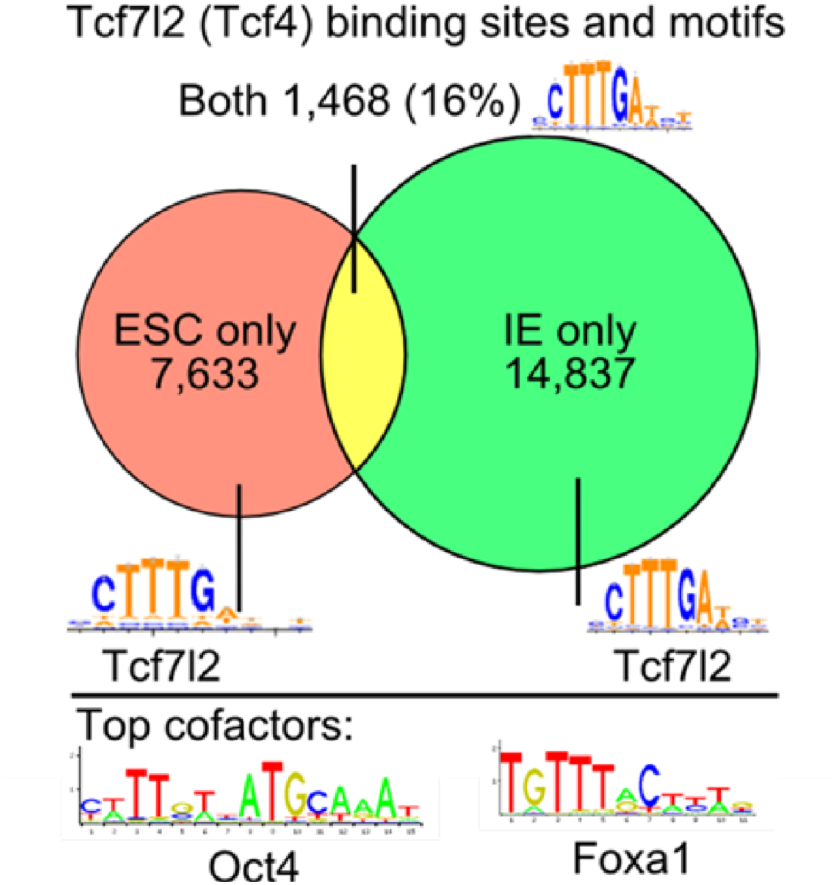
Overlap in binding sites of Tcf7l1 (direct antibody) in mouse embryonic stem cells with Tcf7l2 (FLAG-tagged) in in-vitro differentiated intestinal endoderm, and most enriched non-Tcf motif in these regions.

### 11.2 Experimental Model and Subject Details

All experiments were done in 129P2/OlaHsd mouse embryonic stem cells (mESC), which were cultured according to previously published protocols [ENCODE Project Consortium, 2012]. mESCs were maintained on gelatin-coated plates feeder-free in mESC media composed of Knockout DMEM (Life Technologies) supplemented with 15% defined fetal bovine serum (FBS) (HyClone), 0.1mM nonessential amino acids (NEAA) (Life Technologies), Glutamax (GM) (Life Technologies), 0.55mM 2 *β*-mercaptoethanol (Sigma), 1X E SGRO LIF (Millipore), 5 nM GSK-3 inhibitor XV and 500 nM UO126. Cells were regularly tested for mycoplasma. The non-homologous end joining pathway was disabled by knocking out two necessary genes (Prkdc and Lig4), along with constitutive activation of Rbbp8, which together increase the rate of homologous recombination [Arbab et al., 2015].

### 11.3 Method Details

#### 11.3.1 Phrase library design

The first phrase library (12,000 170bp) was ordered from CustomArray. Two 20bp primer sites were located at each end. One end also included a short (11bp) barcode followed by another primer site (20bp) for separately amplifying it. 11bp is necessay to have enough unique barcodes for all 12,000 sequences such that they are separated by an edit (levenshtein) distance of at least 3, so that any single error to be automatically corrected. Due to the presence of truncated library fragments, however, this barcode couldn’t uniquely identify fragments and wasn’t used. The remaining 99bp was used to test out possible Tcf7l2 binding sites. Combinations of various binding motifs were generated as follows:

1. Sample 25 (out of 35 possible) combinations of the non-Tcf7l2 and non-pioneer motifs (Hnf4a, Gata3, Foxa1/o1, c-Myc, Oct4, Sox3, Smad3).
2. Each of these generates a combination of with and without a Tcf motif.
3. Each of these generates a combination of no pioneer, Ets, Klf, or both motifs.
4. Sample 3 permutations out of each f these.
5. Sample 3 different gap lengths for each sequence.
6. Sample 3 randomly generated sequences for the gap.

The second oligo pool (2,000 149bp) was ordered from Twist Biosciences. Two 25bp primer sites are located at each end, each containing a GATC, flanking a 99bp variable region. Backbones were chosen from the first library based on the probability that the posterior distribution of each phrase’s methylation rate was either lower than, contained, or higher than the average methylation rate of the library (p < 0.05). A set of 30 phrases per backbone were generated by rollong the Tcf motif (ATCAAAG) every 3bp from the starting position. Motifs were scrambled to a specific sequence as follows:

- Tcf: ATCAAAG -> AACGTCG
- Oct: ATGCAAAT -> ATCGGCAT
- Klf: GCCACACCCA -> GCGAGACGCA
- Sox: CWTTGT -> CGTACT
- Gata: AGATA -> ACCCA

**Supplementary Figure 2:**
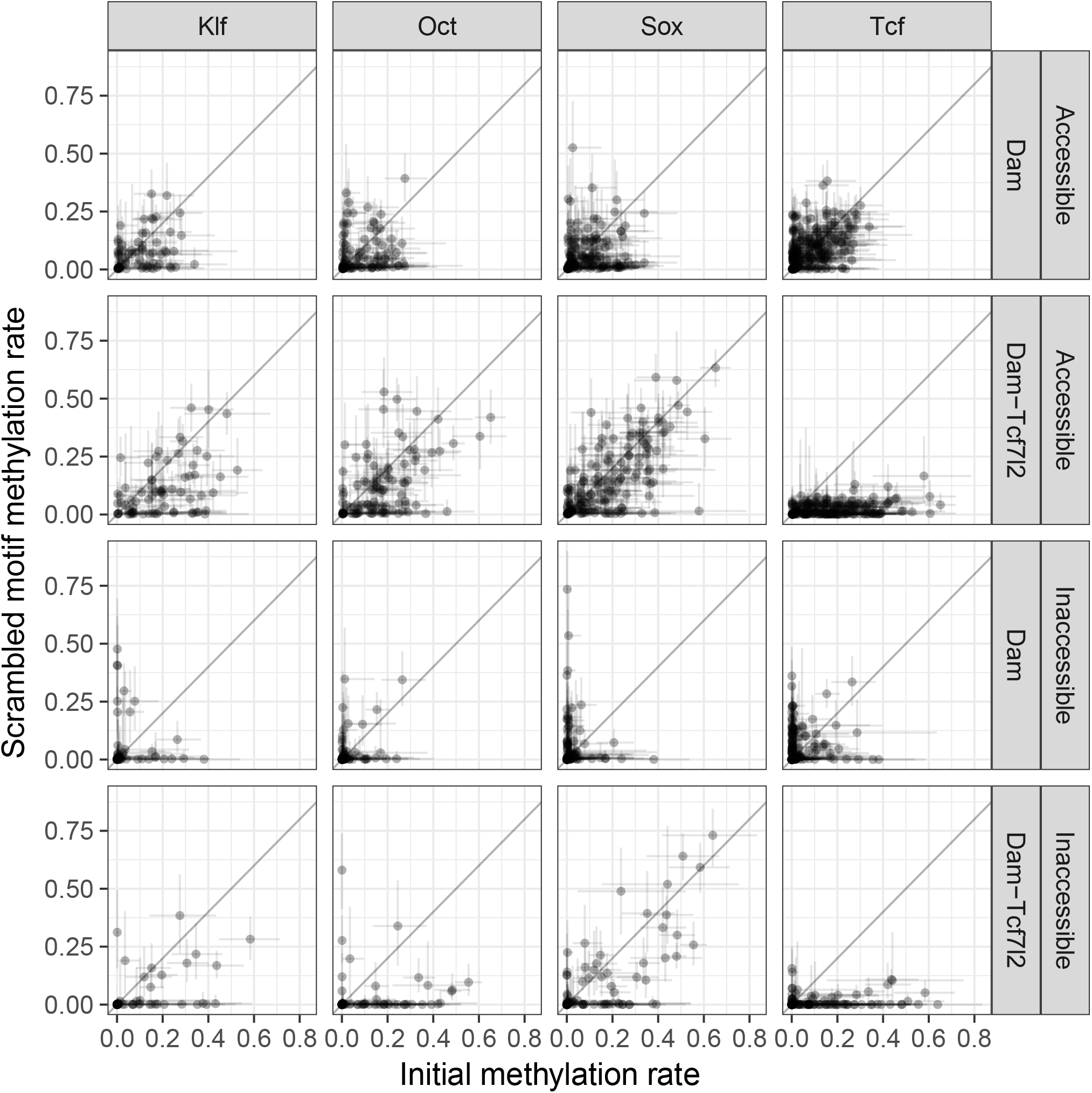
Methylation rate of original phrase and version with scrambled motif. Error bars show the 95% credible interval.

**Supplementary Figure 3:**
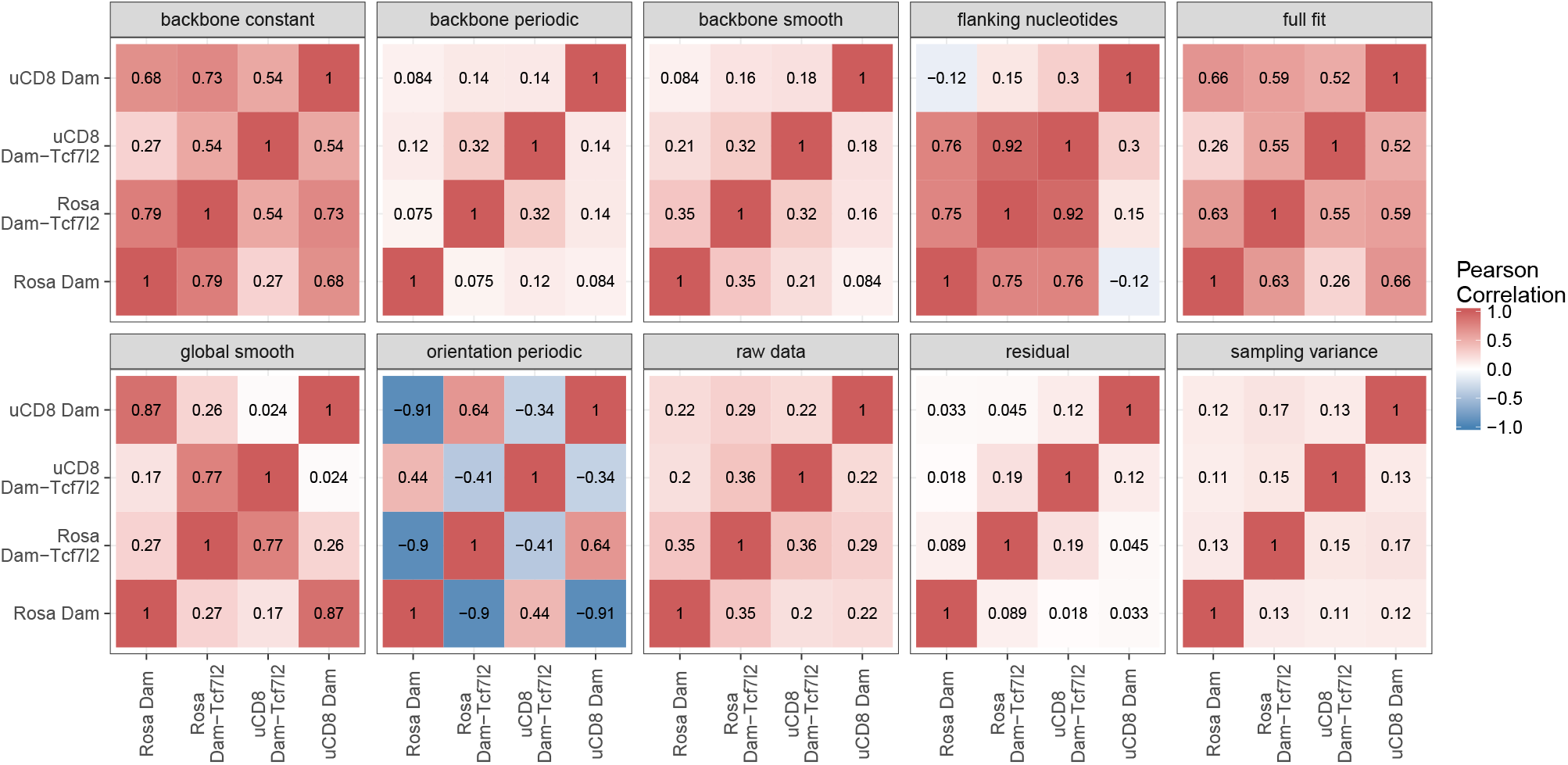
Pearson correlation between the estimated components of each methylation signal between loci and Tcf7l2 fused or unfused Dam. Note: correlation does not capture the magnitude of each signal.

**Supplementary Figure 4:**
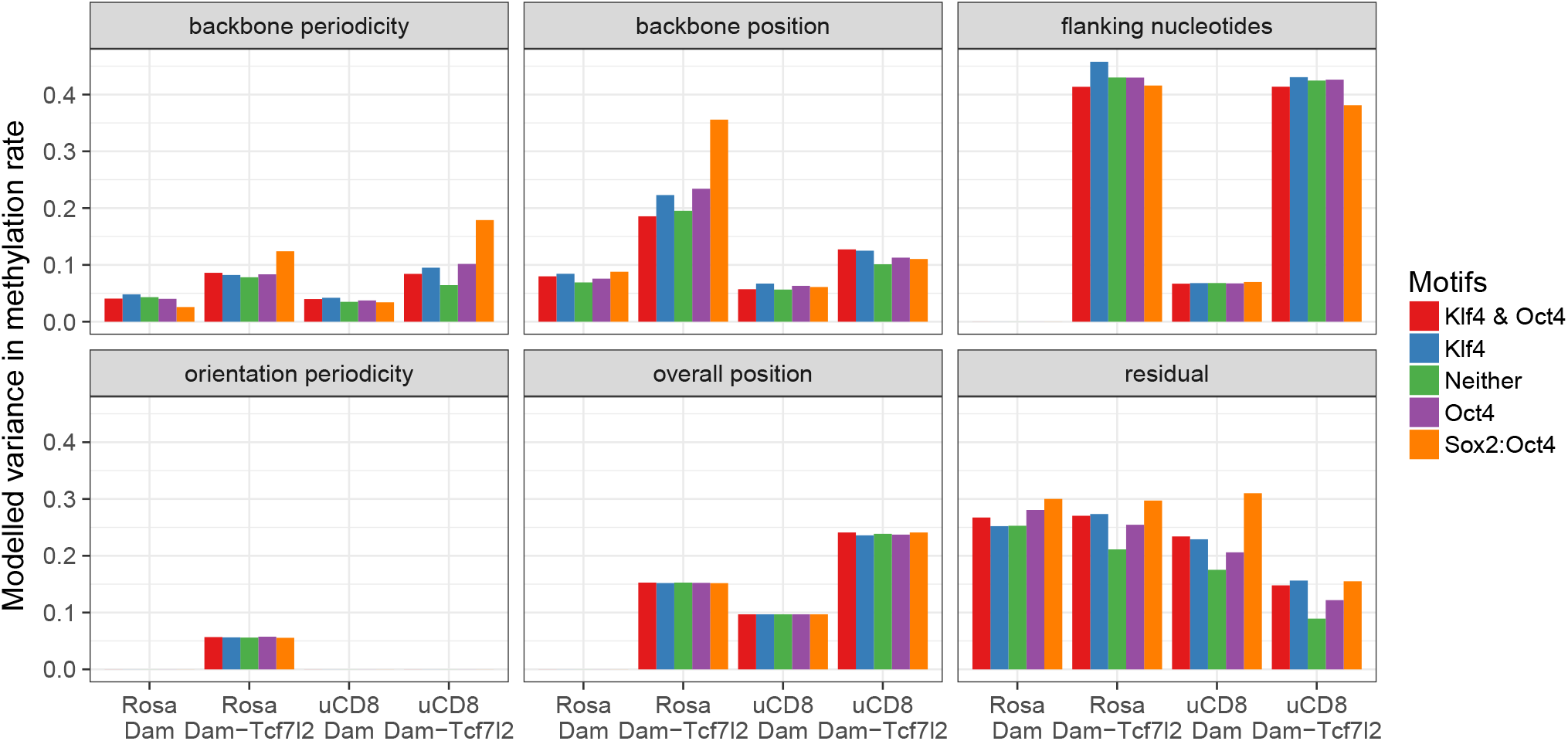
Intra-backbone variance explained by gaussian process components, split into groups of backbones sharing the same cofactor motifs.

**Supplementary Figure 5:**
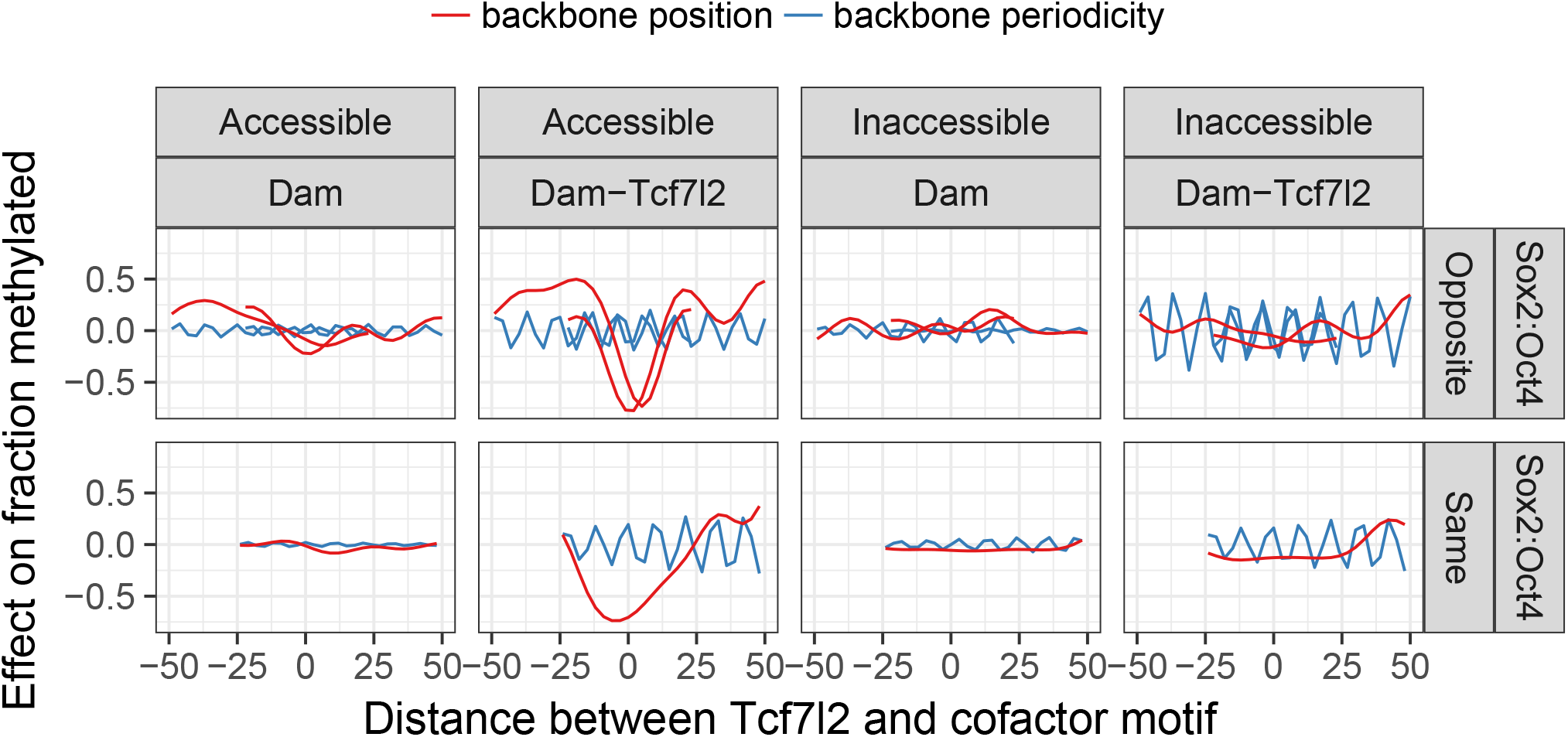
Backbone specifc smooth and periodic components for Dam-Tcf7l2 or unfused Dam methylation for the Sox-Oct containing backbones. Phrases are centered at midpoint of the respective cofactor motif, and calculated to the midpoint of the Tcf motif for each of the possible orientations.

#### 11.3.2 Phrase library integration

Oligo pools were amplified with primers at both ends (40ul NEBnext, 0.2 ul library, Ta=65, 30 cycles) and the 170bp band purified on a 4% agarose gel (Qiagen gel purification). Fragments were extended with homology arms (Table 1) (1000ul NEBnext, Ta = 65, 30 cycles) and purified (Qiagen minelute) to prepare for electroporation. 20 ug CBh Cas9-BlastR plasmid, 20 ug U6-gRNA-HygroR plasmid, and purified fragments were vacuum centrifuged to a final volume of <20 ul, added to 120 ml EmbryoMax Electroporation Buffer (ES-003-D, Millipore), and mixed with mESCs pelleted from a 15cm plate. This was transferred into a 0.4-cm electroporation cuvette and electroporated using a BioRad electroporator (230 V, 0.500 mF, and maximum resistance). Cells were passaged three times following integration.

**Table 1:**
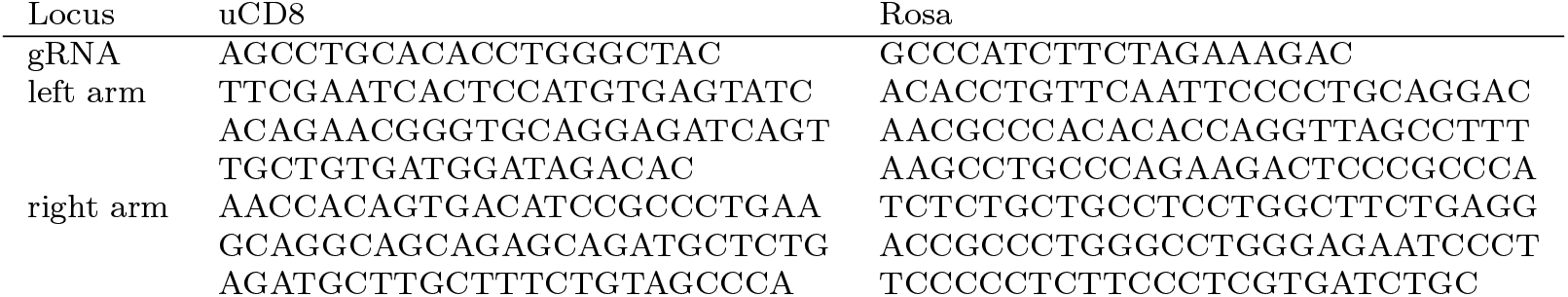
Sequence of guide RNAs and homology arms for CRISPR/Cas9 based homologous recombination.

#### 11.3.3 DamID

Constructs were made by fusing Dam or Dam-N126A to the N-terminus of Tcf7l2 with a short flexible linker [ Szczesnik et a l., 2019]. Dam-Tcf7l2 fusion and Dam only constructs are expressed from a single-integration Dox-inducible transgene expression cassette [Iacovino et al., 2011]. This puts the Dam constructs under control of a tet-responsive promoter, along with integrating a neomycin resistance gene that is selected for by culturing the cells in G418 (300*μ*g/mL) for one week. Following expression of Dam fusion protein (8 hours for wild-type, 24 hours for N126A), genomic DNA is extracted and split into two pools that are digested by either DpnI, which cuts all methylated fragments, or DpnII, which cuts all unmethylated fragments (16ug DNA in 100ul and 100U enzyme).

#### 11.3.4 Next-gen sequencing

Fragments were PCR amplified following DpnI / DpnII digestion with primers outside the homology a rms and spanning the GATC site (16 cycles). This makes integrated fragments at least 100 times more numerous than unintegrated fragments (measured by qPCR), so that they dominate the signal. Two further short PCRs to extend each fragments with adapters for illumina sequencing and a unique barcode for each sample. The resulting fragments are directly sequenced on a next-seq with midoutput 300bp kit (150bp read one, 150bp read two).

### 11.4 Quantification and Statistical Analysis

#### 11.4.1 Alignment

Reads from different samples were demuliplxed with fastq-multx [Aronesty, 2011] based on short barcodes incorporated at the start and end of each sequence. Prior to alignment, overlapping paired end reads were assembled into a single read using PEAR (default parameters) [Zhang et al., 2013], in order to reduce the false positive rate stemming from truncated phrases. The assembled reads are aligned to the ordered library of sequences using BWA (mem algorithm with default parameters) [Li and Durbin, 2009]. Counts are generated for each sequence by summing up all exact matches to it.

#### 11.4.2 Deduplication

A negative-binomial dropout model is used to estimate the amount of amplification, which disambiguates between the situation where most reads are present at high counts due to good coverage from those with poor coverage but high amplificiation. The initial distribution of counts before amplification ( x) for each digest as a negative binomial with shape and rate parameters *α* and *β*. *β* is contrained to be identical between the DpnI and DpnII digests, which prevents the model from assuming different methylation rates between DpnI and DpnII samples. Since phrases are of the same length and similar GC content the observed counts (y) are modelled as arising from a linear amplification of the initial counts: a poisson with rate *γ* * x, where *γ* is the rate of amplification a nd s equencing(Equation 1). This leaves any sequences missing in the original distribution as 0, while 1 shifts to a peak centered on *γ*, 2 to 2*γ*, and so on, creating staggered peaks that eventually run into one another. The dropout reads follow the original unamplified backgorund distribution, rather than a low rate poisson, since it captures the shape of the low level contamination from unintegrated background fragments better in the more lowly amplified samples. Highly amplified samples contain a clear unintegrated fragment population which is amplified in the later library preparation PCR’s. In this case we use a mixture model (R package mixtools) to remove those counts prior to estimating the amplification rate.

To avoid the expensive sum over all possible discrete counts, and since the important information comes from 0 and 1 counts, we approximate higher counts with a continuous distribution. We rewrite our original negative binomial as a poisson-gamma mixture with an explicit latent count rate 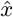, shift the amplification rate to it and simplify down (Equation 2). To avoid double counting the latent genomic counts between 0 and 1 we subtract away which of these continous counts would have arisen from these 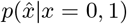 (Equation 3). Equation 4 shows the final likelihood which combines the discrete low counts with continous higher counts, with the structure:

1. Likelihood of unamplified distribution coming from zero counts.
2. Counts from amplified discrete distribution.
3. Counts from amplified continuous distribution.
4. Subtract component of the continous distribution that is already modelled discretely.

*α*, *β*, and *γ* are estimated using hamiltonian monte carlo sampling implemented in the statistical programming language Stan. Since the distribution after normalisation also appears as such, they are treated as iid for further inference and per fragment counts summed up for the respective DpnI and DpnII digests across replicates.

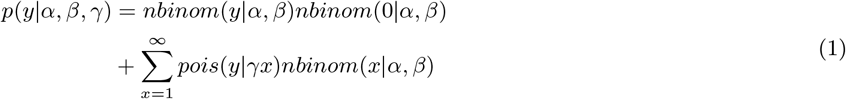

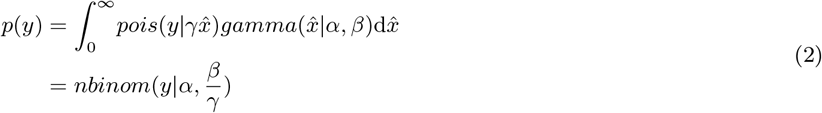

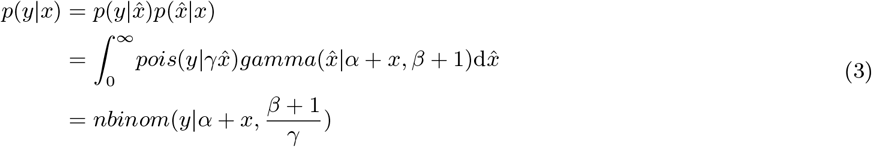

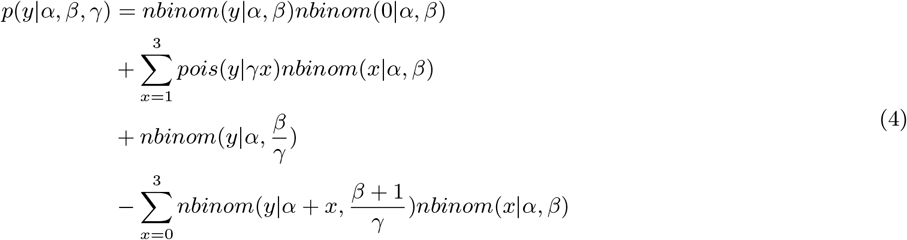

#### 11.4.3 Beta-binomial model

A betabinomial was fit to the normalised DpnI / DpnII count data using the dbetabinom (VGAM package) and mle2 functions (bbmle package) in R. Starting parameters were *α* = *β* = 2 (mean of 0.5 with large spread). Conjugancy the binomial distribution (for read counts) and beta distribution (for methylation rates) results in a straightforward calculation for the posterior beta distribution of methylation rates for each fragment (Equation 5).

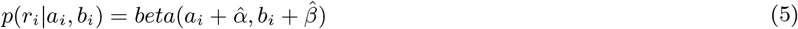

#### 11.4.4 Linear model

Effects of motifs or genomic context were calculated using a generalised linear model (R, glmnet package [Friedman et al., 2010]) with binomial output based on the methylated (DpnII) and unmethylated (DpnI) counts (i.e. logistic regression with multiple measures per point). The regularisation term is optimised via cross-validation.

#### 11.4.5 Gaussian process regression

Gausisan process regression is used to model the methylation rate for each sequence (for an overview of their use in machine learning see [Rasmussen and Williams, 2006]). A logit-normal approximation to the posterior beta function for each sequence (i) was used (Equation 6), where *ψ* and *ψ*_1_ are the digamma and trigamma functions.

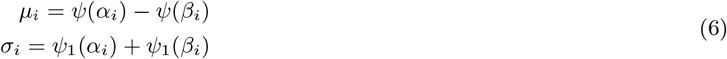

The gaussian process’ behaviour is controlled by the covariance function. Since covariance functions are closed under addition, we combine separate effects by summing up the respective covariance components.

1. Posterior beta variance.
2. Constant effects.
3. Per backbone effects.
4. Per sequence effects.
5. Global positional effects.
6. Orientation positional effects.
7. Backbone positional effects.

Except for the posterior beta variance, which is determined a priori from the data, the remaining components are mixed with separate weights (*w_s_*).

The positional effects are split into two components. A smoothly varying component modelled with a radial basis covariance function, parametrised by a length scale *l*_1_ (Equation 7). A periodic component modelled with a periodic covariance function parametrised by periodicity (*τ*) and length scale (*l*_2_) (Equation 8).

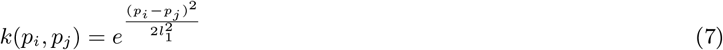

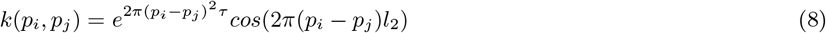

The fit of the model is evaluated by the likelihood fit to the data (Equation 9). The covariance matrix K is constructed by evaluating the covariance function for all pairs of sequences (with residuals adding in as the extra identity matrix). Cholesky decomposition is used for inverting the covariance matrix.

Different components of the gaussian process fit are extrated using Equation 10, where the K that is inverted can be varied to include only the relevant portions of the covariance function. Cross validated fits are similarly calculated as in Equation 11 based on a formula for linear smoothers [Cook and Weisberg, 1982].

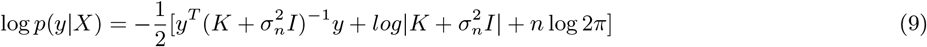

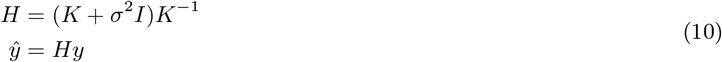

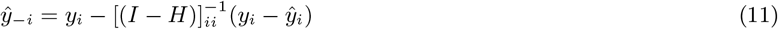

The weights of the various components, and the hyperparameters (legnth and periodicity) were found by gradient based optimisation of the likelihood using L-BFGS. The opimal periodicity was found by grid search over all integer periodicities (up to 30bp) prior to gradient based optimisation. All algorithms in this section in Haskell through bindings to linear algebra (Blas and Lapack) and optimisation (libLBFGS) libraries.

